# The genetic diversity of honeybee colonies predicts the gut bacterial diversity of individual colony members

**DOI:** 10.1101/2021.06.04.447042

**Authors:** C Bridson, L Vellaniparambil, R E Antwis, W Müller, R T Gilman, J K Rowntree

**Author notes:** Corresponding author: Dr Jennifer K Rowntree.

## Abstract

1. The gut microbiota of social bees is relatively simple and dominated by a core set of taxa that have been reported consistently in individual workers from around the world. Yet, variation remains, and this has been shown to affect host health.
2. We characterised the individual- and regional-scale variation in the honeybee (*Apis mellifera*) gut microbiota in the North West of England, and asked whether the microbiota was influenced by host genotype or landscape composition.
3. We collected multiple honeybees from 64 colonies, and sequenced the V4 region of the 16S rRNA gene to characterise the mid- and hindgut bacterial communities. We characterised the genotype of each individual honeybee, and also the land cover surrounding each colony.
4. The literature-defined core taxa consistently dominated across the region, despite the varied environments. However, there was variation in the relative abundance of core taxa, and colony membership explained a large proportion of this variation. Individuals from more genetically diverse colonies had more diverse microbiotas, but individual genetic diversity did not influence gut microbial diversity. There was a trend for colonies in more similar landscapes to have more similar microbiota, whilst bees from more urban landscapes had a slightly less diverse microbiota than those from less urban landscapes.
5. Our study provides, to our knowledge, the first demonstration for any species that the gut bacterial communities of individuals can be influenced by the genotypes of other conspecifics in the population. This is particularly important for social organisms, such as honeybees, as colony rather than individual genetic diversity appears to drive gut microbial diversity, a factor related to colony health.

## Introduction

Microbial symbionts are ubiquitous in their association with animals, and their influence on host species ranges across a spectrum from deleterious to beneficial (Ferrari and Vavre, 2011). An important class of symbionts resides in the gut. Gut microbial communities can affect host health (Kühn *et al*., 1993; Round and Mazmanian, 2009; Sekirov *et al*., 2010; Clemente *et al*., 2012), ecology, and evolution (Dillon and Dillon, 2004; Brucker and Bordenstein, 2012; Engel and Moran, 2013; McFall-Ngai *et al*., 2013; Rosenberg and Zilber-Rosenberg, 2016), for example by influencing host behaviour (Cryan and Dinan, 2012), protecting against parasites (Koch and Schmid-Hempel, 2011a), or providing insecticide resistance (Kikuchi *et al*., 2012).

Social bees provide a good model system for studying gut microbiota (Zheng *et al*., 2018). The bee gut microbiota is relatively simple, and is dominated by a core of nine bacterial phylotypes that are found consistently across continents, genotypes, and landscapes (Jeyaprakash *et al*., 2003; Mohr and Tebbe, 2006; Babendreier *et al*., 2007; Cox-Foster *et al*., 2007; Ahn *et al*., 2012; Disayathanoowat *et al*., 2012; Martinson *et al*., 2012; Moran *et al*., 2012; Sabree and Moran, 2014; Corby-Harris *et al*., 2014a; Kwong *et al*., 2017). These phylotypes include six Proteobacteria [*Giliamella apicola, Snodgrassella alvi* (Kwong and Moran, 2013)*, Frischella perrara* (Engel *et al*., 2013a)*, Bartonella apis* (Kešnerová *et al*., 2016), Alpha 2.1 (*Commensalibacter*), and Alpha 2.2 (*Parasaccharibacter apium*) (Martinson *et al*., 2011; Corby-Harris *et al*., 2014b; Corby-Harris and Anderson, 2018)], two Firmicutes [*Bombilactobacillus* spp. (formerly *Lactobacillus* Firm-4) and *Lactobacillus* Firm-5 (Killer *et al*., 2014; Olofsson *et al*., 2014; Zheng *et al*., 2020)], and an Actinobacterium [*Bifidobacteria asteroides* (Scardovi and Trovatelli, 1969; Bottacini *et al*., 2012; Ellegaard *et al*., 2015)].

Despite the consistent presence of these phylotypes in the bee gut bacterial community, there is variation in the relative abundance of the core phylotypes and in the core strains present (Moran *et al*, 2012; Engel *et al*., 2012; Ellegaard and Engel, 2016; Kwong and Moran, 2016; Powell *et al*., 2016). Variation in bacterial composition can lead to functional differences (Engel *et al*., 2012) in traits such as pollen and saccharide breakdown (Engel *et al*., 2012; Lee *et al*., 2015) and resistance to disease (Koch and Schmid-Hempel, 2012). Until now studies have focused on the differences among regions and continents, but there is little knowledge of how the bee microbiota composition varies within regions that comprise multiple apiaries across a landscape. At this scale, the set of apiaries can be considered a population or metapopulation, with the potential for genetic material to be transferred among them. Understanding variation in the microbiota at the regional level is important because this variation can influence how host populations respond to change and their resilience to parasites and pesticides. (Zilber-Rosenberg and Rosenberg, 2008; Hehemann *et al*., 2012; Koch and Schmid-Hempel, 2012; Quercia *et al*., 2014; Suzuki, 2017). Furthermore, if individual colonies have distinct microbiotas, then this may drive differences among colonies in fitness and health, and thus could be important for understanding and managing bee declines or optimising honey production (Ribière *et al*., 2019).

Studying multiple colonies across a region can help us to understand the variables that influence the honeybee microbiota, for example the effect of different land use types on the honeybee gut microbial community (Engel *et al*., 2016). Land use can be a proxy for floral composition (Kleijn *et al*., 2006), and potentially for other environmental factors such as exposure to pollutants (Botías *et al*., 2017). Land use change is also thought to be a major factor in bee decline (Potts *et al*., 2010), and land use per se has been shown to influence gut microbiota composition in other systems (Teyssier *et al*., 2018), yet there is little knowledge of how it affects bee microbiota composition. Host genotype has also been shown to shape host microbiotas in some systems (Zoetendal *et al*., 2001; Bevins and Salzman, 2011; McKnite *et al*., 2012; Goodrich *et al*., 2014a; Griffiths *et al*., 2019). This may be particularly important for honeybee colonies because the majority of workers in a colony are related to the queen, although as honeybee queens are promiscuous, workers are not always full sibs (Estoup *et al*., 1994).

In this study, gut bacterial communities of multiple individuals were sampled from 64 honeybee colonies across the North West of England, UK, providing high landscape coverage. There were four main aims of the study: (i) to characterise the gut bacterial community composition of honeybees in the North West of England; (ii) to investigate how variation in the bacterial community is partitioned among colonies in the region; and (iii) to determine whether landscape diversity and composition and/or (iv) the genetic diversity of colonies or individuals influence the honeybee gut bacterial community.

## Materials and methods

### Sample collection

We sampled individual honeybees from 64 colonies across North West England (Fig. 1) in May and June 2016. The colonies were spread across both rural and urban environments, including three major urban centres, and were mostly owned by amateur beekeepers who volunteered to participate in the study. We sampled one colony per apiary, and five randomly selected nurse bees per colony. Nurse bees are adult workers that have yet to leave the nest, and that perform roles such as feeding the brood and guarding the nest (Winston, 1987). We sampled the nurse stage because these bees take food from multiple foragers and distribute it to the larvae. Hence, they are exposed to a wide variety of food sources, and their microbiota may better reflect that of the colony as a whole (Kapheim *et al*., 2015). We identified nurse bees by shaking a brood frame into a bucket and collecting only those bees that did not fly away. After collection, bees were euthanised in 100% ethanol and frozen at -20 °C until further use (Koch and Schmid-Hempel, 2011b; Powell *et al*., 2014).

**Figure 1.**
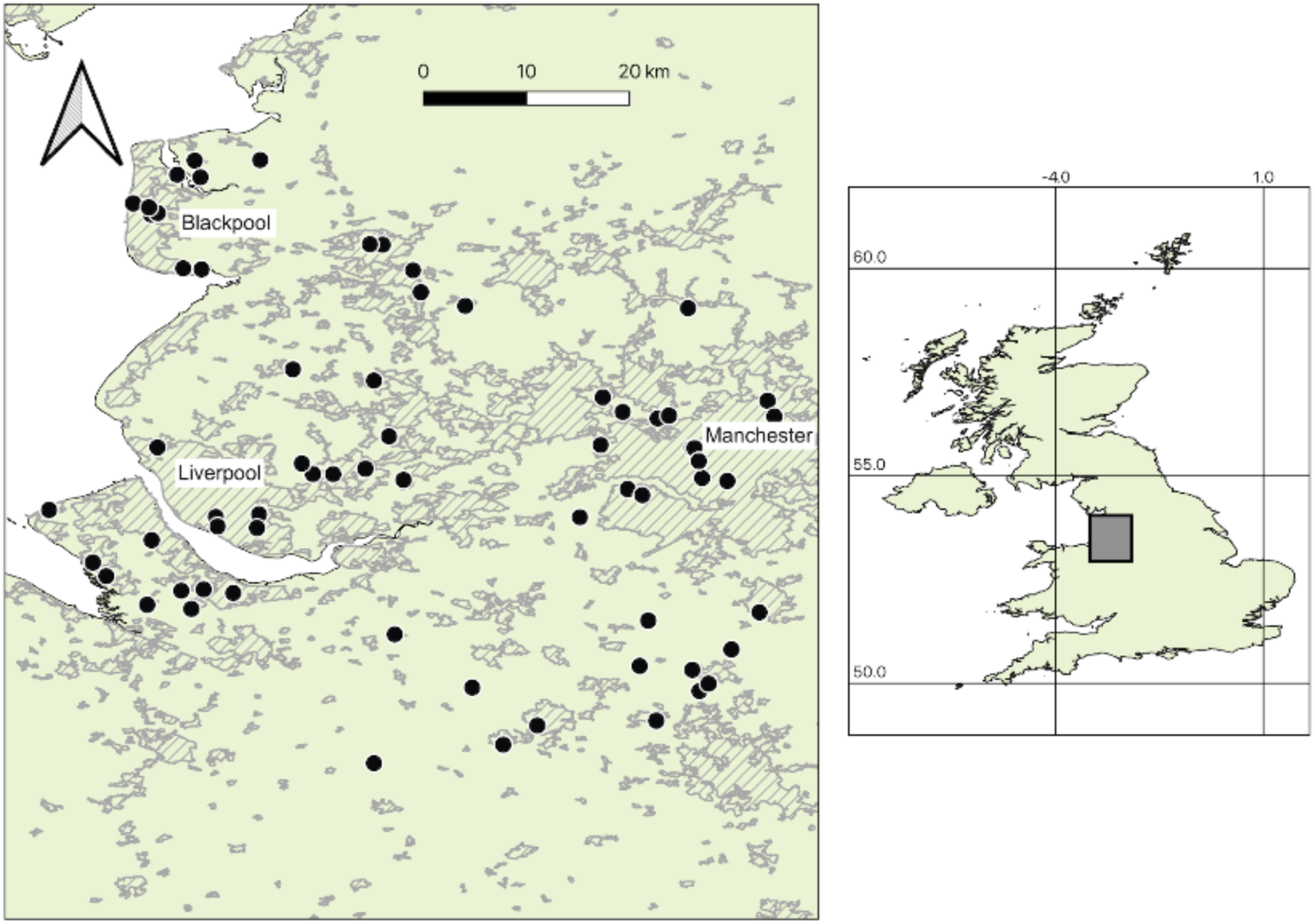
Map of the study area in North West England with location shown as a grey box on the inset map of the UK. Colonies are shown as black dots and urban areas as hatched grey. The UK layer was downloaded from map.igismap.com and the urban layer (showing built areas in December 2011) from geoportal1-ons.opendata.arcgis.com.

### Dissection and DNA extraction

Bees were washed in 1% (v/v) sodium hypochlorite solution for five minutes, followed by three washes in autoclaved MilliQ water, to remove any external bacteria that could contaminate the sample. To further reduce contamination, we carried out dissection, DNA extraction, and other molecular procedures in a PCR hood (UVP, CA) in which all equipment and surfaces had been UV-sterilised for 30 minutes. Dissecting instruments were autoclaved before use, and then cleaned with 1% sodium hypochlorite solution and heat treated to 250 °C using a bead steriliser (Steri 250, Inotech, UK) between dissections of bees from different colonies. Dissection involved removing the head, legs and wings, then using micro-dissecting Vannas spring scissors (FST, Germany) to cut a window into the dorsal cuticle. We teased the ventriculus away from the body wall using a needle, and used forceps to pull out the gut tract. If the crop did not separate naturally, it was cut away. Only the midgut and hindgut were used for this study, with one gut tract used per sample. The process was carried out without a microscope to reduce the possibility of contamination. We carried out negative extractions each day to monitor any contamination, which equated to two or three colonies per negative extraction. All negative extraction samples were processed as normal, but instead of adding honeybee guts, sterilised forceps were scraped against the inside of the Eppendorf.

We placed dissected guts in sterile 2-ml Eppendorfs with 180 μl SDS lysis buffer (200 mM NaCl, 200 mM Tris, 20 mM EDTA, 6% SDS; Moran *et al*., 2012). To homogenise the guts, we added a 5-mm stainless steel bead (Qiagen, UK) and 500 μl of 0.1 mm acid-washed glass beads (Sigma-Aldrich, UK) to the sample, then shook the sample twice for 2.5 minutes at 30 Hz on a Mixer Mill, inverting samples between shakings (Retsch, Germany; Engel *et al*., 2013b). After bead-beating, we added of 30 μl of 10 mg/ml lysozyme (Sigma-Aldrich, UK) and incubated the samples at 37 °C for 1 hour to digest Gram-positive cell walls (Teng *et al*., 2018). We extracted DNA using the Qiagen DNeasy Blood and Tissue Kit following the protocol for Gram-positive bacteria from step 4 of the DNeasy handbook (Qiagen DNeasy Blood and Tissue Handbook, 2006). The only modification to this protocol was to incubate the samples in proteinase K and buffer AL for 1 hour at 56 °C after the lysozyme step. The DNA was eluted in 200 μl of Buffer AE and stored at -20 °C until further use.

### Sequencing the gut bacterial community

We sequenced the gut bacterial communities using a modification of the dual-index sequencing protocol described in Kozich *et al*. (2013) and Antwis *et al*. (2018). Briefly, the hypervariable V4 region of the 16S rRNA gene was amplified using the universal primers 515F (TATGGTAATTGTGTGCCAGCMGCCGCGGTAA) and 806R (AGTCAGTCAGCCGGACTACHVGGGTWTCTAAT), modified with Illumina adaptors and index sequences. The V4 region is commonly used to analyse bacterial communities and has previously been used to study the bee microbiota (Powell *et al*., 2014; Raymann *et al*., 2017; Zheng *et al*., 2017). The V4 region was amplified in triplicate for each sample in a single PCR step using the following conditions: 95 °C for 15 min; 25 cycles of 95 °C for 20 s, 55 °C for 60 s, 72 °C for 60 s; and then 72 °C for 10 min. The triplicate PCRs were pooled, and the amplicons were cleaned using Agencourt Ampure beads (Beckman Coulter Genomics, Indianapolis, IN, USA) and quantified and quality checked using a Tapestation (Agilent Technologies Inc., CA, USA). DNA concentrations were determined using Qubit (Thermo Fisher Scientific, USA). We removed 29 samples because they had insufficient concentrations of DNA, leaving two to five samples (i.e., bees) per colony. Equimolar amounts of each remaining sample were pooled and sequenced together, along with 10% PhiX v3 (Illumina, San Diego, CA, USA) as an in-run control. Sequencing was conducted using Illumina v2 chemistry (2×250bp paired-end sequencing) on the MiSeq platform (Illumina, San Diego, CA, USA) at the University of Salford.

### Bioinformatics

We processed sequence data using the Dada2 v1.6.0 pipeline (Callahan *et al*., 2016) with default parameters in R 3.4.1 (R Core Team, 2017). Dada2 has been shown to perform well in relation to other pipelines (Callahan *et al*., 2016; Nearing *et al*., 2018). We trimmed the raw sequences when the Phred score (Q) was less than 30, which was at 240 bp for the forward read and 230 bp for the reverse. Using the ‘filterandTrim’ function, we truncated reads when the Illumina quality score was less than two (truncQ = 2), and we removed sequences when the maximum expected error (maxEE) was greater than two. The modal contig length of the merged paired-end reads was 233 bp. Of the 1279 total amplicon sequence variants (ASVs) obtained, we removed 36 for having lengths greater than 236 bp, and we removed 883 chimeras using the function ‘removeBimeraDenovo’. Taxonomic assignment of the remaining sequences was performed using the RDP Naïve Bayesian Classifier method described in Wang *et al*. (2007), with training sets constructed from the Silva v128 database (Quast *et al*., 2012) using Dada2’s default parameters.

We removed ASVs that were not taxonomically assigned to the kingdom ‘Bacteria’ or that were classified to the class ‘Chloroplast’ or the family ‘Mitochondria’. At this point, all remaining ASVs were assigned down to at least the phylum level. We removed ASVs that were only present in one sample, as these are more likely to be erroneous (Goodrich *et al*., 2014b). The 52 negative controls contained very low levels of contamination, with the total reads in each negative control ranging from 7 to 118. Across the negative controls, there were 18 ASVs that were also found in the true samples. We removed ASVs if the mean number of reads across all true samples was not at least 10x higher than in the negative controls. This led to the removal of seven ASVs, all of which were known contaminants from previous studies (Salter *et al*., 2014). The other 11 ASVs in the negative controls were all members of the previously-identified core phylotypes, and were the most abundant ASVs found in this study. After all filtering, 136 of the original 1279 ASVs remained in our analyses. We generated rarefaction curves for each sample using the command ‘rarecurve’ in the package vegan (Oksanen *et al*., 2018; Supplementary Figure S1).

We applied maximum likelihood analysis to construct a phylogenetic tree of the filtered ASVs using the phangorn package v2.3.1 (Schliep, 2011) with a GTR+G+I nucleotide substitution model. We chose the GTR+G+I nucleotide substitution model because it had the highest log likelihood and lowest Akaike Information Criterion (AIC) and Bayesian Information Criterion (BIC) among available model options (Posada & Buckley, 2004). The tree was midpoint rooted using the phytools package (Revell, 2012).

### Measuring biodiversity of the gut bacterial community

We calculated beta diversity from the ASV abundance data. Beta diversity captures variation among ecological communities (i.e., how gut bacterial communities differ among bees or colonies). Due to unequal sequencing depth across samples, we used Cumulative Sum Scaling (CSS; Paulson *et al*., 2013) to produce a normalised matrix of ASV abundances. Results obtained using CSS-normalised data were consistent with those obtained using data normalised by variance stabilisation transformation in the DESeq2 package (Love *et al.,* 2014). We generated CSS-normalised ASV abundance matrices for both individual bees and colonies. Abundance at the colony level was calculated by summing the normalised abundance values for each ASV across all samples from a colony and then dividing by the number of samples from that colony. We used weighted UniFrac distances to produce pairwise dissimilarity matrices from the CSS-normalised abundance data at the individual and colony levels by applying the ‘distance’ function in phyloseq 1.21.0 (McMurdie and Holmes, 2013). UniFrac distances measure dissimilarity accounting for phylogeny (Lozupone *et al*., 2011). The dissimilarity matrices were visualised with non-metric multidimensional scaling (nMDS) performed by the ‘ordinate’ function in phyloseq.

We measured alpha diversity using Hill numbers with q values of 0, 1 and 2, which differ in how they weight rare ASVs (Hill, 1973). Alpha diversity captures the diversity of the bacterial community within individual bees or colonies. When q = 0 the Hill number reports absolute species richness, and when q > 0 the Hill number increases as the combination of richness and evenness of the community increases. For individual bees, we calculated Hill numbers from the raw read counts using rarefaction/extrapolation curves to account for differences in sequencing depth among samples (Chao *et al*., 2014). For colonies, we estimated Hill numbers from the mean proportional abundance of ASVs across all samples from the colony. Thus, the Hill numbers for colonies weight each sample from the colony equally regardless of sampling depth, but are sensitive to the number of samples from the colony.

### Landscape analysis

We determined the landscape composition surrounding each colony (except one for which the GPS location was not recorded) using data from the Land Cover Map 2015 (25m raster, GB; Rowland *et al*., 2017). The Land Cover Map classifies the land cover of each 25 m^2^ pixel into one of 21 land cover classes. We used QGIS version 2.18.14 (QGIS Development Team, 2017) to overlay the Land Cover Map raster layer onto the locations of colonies, and to define buffer zones around each colony at 500 m, 1.5 km, 5 km and 10 km radii. Buffer zone sizes were chosen based on the frequency at which honeybees forage at different distances from the colony (Visscher and Seeley, 1982; Waddington *et al*., 1994; Steffan-Dewenter and Kuhn, 2003). Honeybees can forage up to 10km from the colony (Visscher and Seeley, 1982), whilst the 500m buffer zone was included to investigate local colony effects. The 1.5 km buffer zone was chosen to reflect the mean foraging distance reported in a previous study (Steffan-Dewenter and Kuhn, 2003), whilst 90% of honeybees have been shown to forage within 5 km (Visscher and Seeley, 1982). We used the LecoS plugin (Jung, 2016) for QGIS to determine the proportion of each buffer zone that belonged to each of the 21 land cover classes, which includes one urban class. The landscape diversity of each buffer zone around each colony was determined by estimating the Hill number (q = 1) from the raw land-use proportion data using the package hillR (Li, 2018). The suitability of Hill numbers to estimate landscape diversity is supported by the fact that the related Shannon index is widely used to characterise landscape diversity (Nagendra, 2002; Ramezani, 2012).

### Genotyping bees and colonies

We determined individual bee genotypes at five microsatellite loci according to the protocol described in Evans *et al*. (2013) using the primers detailed in Supplementary Table S1. These microsatellite loci have been used in other honeybee population genetic studies (Muñoz *et al*., 2009; Muñoz *et al*., 2012). We ran the microsatellites as two separate multiplexes, with primers A113, AP043, and AP055 in multiplex 1 and primers A007 and B124 in multiplex 2, using the Qiagen Type-It master mix kit (Qiagen, Valencia, CA) in a final reaction volume of 10 µl. PCR amplification involved denaturation at 94 °C for 5 min, followed by 30 cycles of 95 °C for 30 s, 57 °C for 30 s, and 72 °C for 30 s, with a final elongation step at 72 °C for 30 min. The size of the microsatellites was determined by capillary electrophoresis on an ABI 3730 DNA Analyser (Applied Biosystems, CA, USA) at the University of Manchester Sequencing Facility, using the GeneScan^TM^ LIZ500 (Thermo Fisher Scientific, USA) size standard. The output peaks were scored using Genemapper v5.0 software (Applied Biosystems, Foster City, CA, USA) and sorted into allele bins using the MsatAllele package v1.05 (Alberto, 2009) in R (R Core Team, 2017). A subset of samples were repeated to ensure consistency of allele calling.

We analysed host genotype data using the package adegenet (Jombart, 2008). We calculated the number of alleles per locus, whether each locus was in Hardy-Weinberg equilibrium, and the expected and observed heterozygosities in the population. To measure the genetic diversity of individual bees, we calculated the heterozygosity of each individual. To measure the genetic diversity of colonies, we calculated i) the mean observed heterozygosity of each colony and ii) the mean proportion of alleles shared between pairs of bees in the colony.

### Statistical analyses

We used our weighted UniFrac dissimilarity matrices to investigate the partitioning of variability in gut bacterial communities within and among colonies, and to ask whether colonies or individuals with more similar landscapes or genotypes also had more similar gut bacterial communities. We used regression analysis to investigate how the alpha diversity of bacterial communities at the colony and individual levels varied over space and as a function of landscape composition and host genetic diversity.

#### How is variation in the gut bacterial community partitioned within and among colonies?

We partitioned variation in the gut bacterial community within and among colonies using permutational multivariate analysis of variance (PERMANOVA; Anderson, 2001) with 9999 permutations applied to our weighted UniFrac dissimilarity matrices and implemented in the ‘adonis’ function of the package vegan (Oksanen *et al*., 2018). We used a K-Nearest Neighbour (KNN) graph-based analysis of the CSS-normalised individual-level data, implemented with the functions ‘phyloseqGraphTest’ and ‘igraph’ (Csárdi and Nepusz, 2006), to ask whether the individuals with the most similar microbiotas were likely to be nestmates or from different colonies. We performed the KNN analysis for K=3, because individuals from most colonies had at least three nestmates that were sampled.

#### Do colonies that are closer in space have more similar bacterial communities?

We studied the relationship between colony bacterial community composition and the geographic distance between colonies using a Mantel test (9999 permutations; Mantel, 1967). The geographic distance matrix was created by converting the longitude and latitude coordinates of each colony into UTM coordinates using the R package rgdal (Bivand *et al.,* 2020).

#### Does land use affect gut bacterial community composition?

We used partial Mantel tests (9999 permutations) to assess whether colonies with more similar surrounding landscapes had more similar bacterial communities when controlling for distance between colonies. We created the bacterial community composition matrix from pooled colony data. Distance matrices for land use composition were generated for each of the four buffer zones using Bray-Curtis dissimilarities calculated with the function ‘vegdist’ in the package vegan (Oksanen *et al*., 2018). We included the geographic distance matrix in the analysis to account for potential spatial autocorrelation in gut bacterial community composition. We performed partial Mantel tests for each buffer zone separately.

Urbanisation has been shown to influence gut microbiota in other systems (Teyssier *et al*., 2018). Therefore, we performed a PERMANOVA using the function ‘adonis’ to ask whether the pooled colony gut bacterial community composition varied with the proportion of urban land surrounding the colony.

#### Do genetic diversity and host genotype affect gut bacterial community composition?

We generated genetic dissimilarity matrices (Roger’s genetic distances; Rogers, 1972) from the individual and colony-level multilocus allele data using the package adegenet (Jombart, 2008), and visualised genetic dissimilarity using nMDS. We used a Mantel test (9999 permutations) to determine whether individuals with more similar genotypes harboured more similar bacterial communities. For the colony-level data, we used a partial Mantel test, which also incorporated the geographic distance matrix, to account for potential spatial autocorrelation in gut bacterial community composition.

#### What factors predict alpha diversity in the gut bacterial community at the colony and individual levels?

To understand the factors that predict the diversity of the gut microbiome at the colony level, we regressed the colony-level Hill numbers on i) the mean proportion of alleles shared between individuals in the colony, ii) the mean heterozygosity of the colony, iii) the landscape diversity in the 5 km buffer zone around the colony, iv) the proportion of urban land surrounding the colony, and the v) northing and vi) easting of each colony. We included the number of bees sampled from the colony as a categorical predictor in the model. This controls for the fact that we expect to find more ASVs in colonies from which more bees were sampled, but makes no assumptions about the rate at which ASVs accumulate as the number of samples increases. We weighted the variance of each colony-level data point in proportion to the square root of the number of samples (i.e., individual bees) from which that data point was calculated. Initially, we fitted linear regressions using the package nlme (Pinheiro *et al*., 2020) which allowed us to include spatially autocorrelated error in the model. However, we found no evidence for spatial autocorrelation, and therefore we removed spatial autocorrelation from the model and fitted linear regressions using the function lm in base R. For each Hill number (i.e., q ∈ {0, 1, 2}), we fitted models that included each possible combination of the predictors in the full model. We weighted each model according to its Akaike weight, and calculated the effect size of each predictor as the weighted average across all models in which that predictor appeared (Burnham & Anderson, 2002). To obtain p-values for each predictor, we computed the confidence distribution around the effect size of the predictor in each model in which it appeared, weighted these according to the Akaike weights of the models, and summed across all models. The p-value associated with the predictor is two times the minimum of the proportion of the confidence distribution that is greater than zero or is less than zero (Gilman *et al.,* 2018).

To understand the factors that predict the diversity of the gut microbiome of individuals within colonies, we regressed the individual-level Hill numbers on the heterozygosity of the individual and on all six colony-level predictors. We included a random effect of colony in the model, and fitted models by maximum likelihood using the package lme4 (Bates *et al.,* 2015). For each Hill number (i.e., q ∈ {0, 1, 2}), we calculated the effect size and p-values associated with each predictor as described for the colony-level analysis.

## Results

A total of 6,900,477 reads passed the quality filter thresholds, from an initial output of 9,487,873 reads from the MiSeq run. The number of reads per sample ranged from 7,758 to 44,343 (mean 23,551), with a total of 136 ASVs detected across the study. The rarefaction curves for each sample reached a plateau (Supplementary Figure S1). Thus, the sequencing depth was sufficient to characterise the microbial community of the samples.

### Honeybee gut bacterial community composition in the North West of England

In line with previous studies, 98.9% of reads were members of the literature-defined core taxa, and only six samples had bacterial community compositions that consisted of less than 90% core phylotypes. All core phylotypes except *Parasaccharibacter apium* were found in this study.

Each core taxon present in this study was represented by multiple ASVs, but the majority of ASVs were rare (Supplementary Table S2). The 20 most abundant ASVs comprised 94.5% of all reads in the study. These included representatives from each of the nine core phylotypes except *P. apium,* and also included the non-core species *Lactobacillus kunkeei*. Thus, the pattern of dominance by the core phylotypes is driven by just a one or two ASVs from each taxon. The most abundant core phylotypes in this region were *Lactobacillus* Firm 5 and *Gilliamella apicola*, which together constituted 48.6% of the regional honeybee gut bacterial community (Supplementary Table S2). Outside of the core, the majority of ASVs belong to the family *Enterobacteriacae*.

All of the core phylotypes found in this study were present in each of the colonies (Supplementary Table S3). In contrast, only 67% of individuals possessed all of the core phylotypes, and 7.8% of individuals were missing two or more phylotypes. However, only the *Commensalibacter* phylotype was present in less than 90% of individuals.

### How is variation in the gut bacterial community partitioned within and among colonies?

Colony membership had a significant effect on honeybee gut bacterial community composition, explaining 41% of the variation among samples (PERMANOVA: R^2^=0.414, df=63, 229, p=0.0001; Fig. 2). There were marginally significant differences in sample variance within colonies (betadisper: F=1.38, p=0.051), but PERMANOVA is robust to differences in variance when designs are well-balanced (Anderson and Walsh, 2013). Due to the differences among colonies in sample size, we randomly selected three individuals from each colony to ensure there was an equal number of observations per colony, and performed the PERMANOVA on this subset of the data, using 9999 permutations as before. We repeated the analysis 50 times, with each analysis producing the same p value. KNN analysis showed that the gut bacteria of bees from the same colony were more similar than expected by chance (p=0.001). However, only 5.7% of edges were between samples from the same colonies. Thus, while there is structure by colony, high variation within colonies means that bees from the same colony did not have distinguishable colony-specific microbiota.

**Figure 2.**
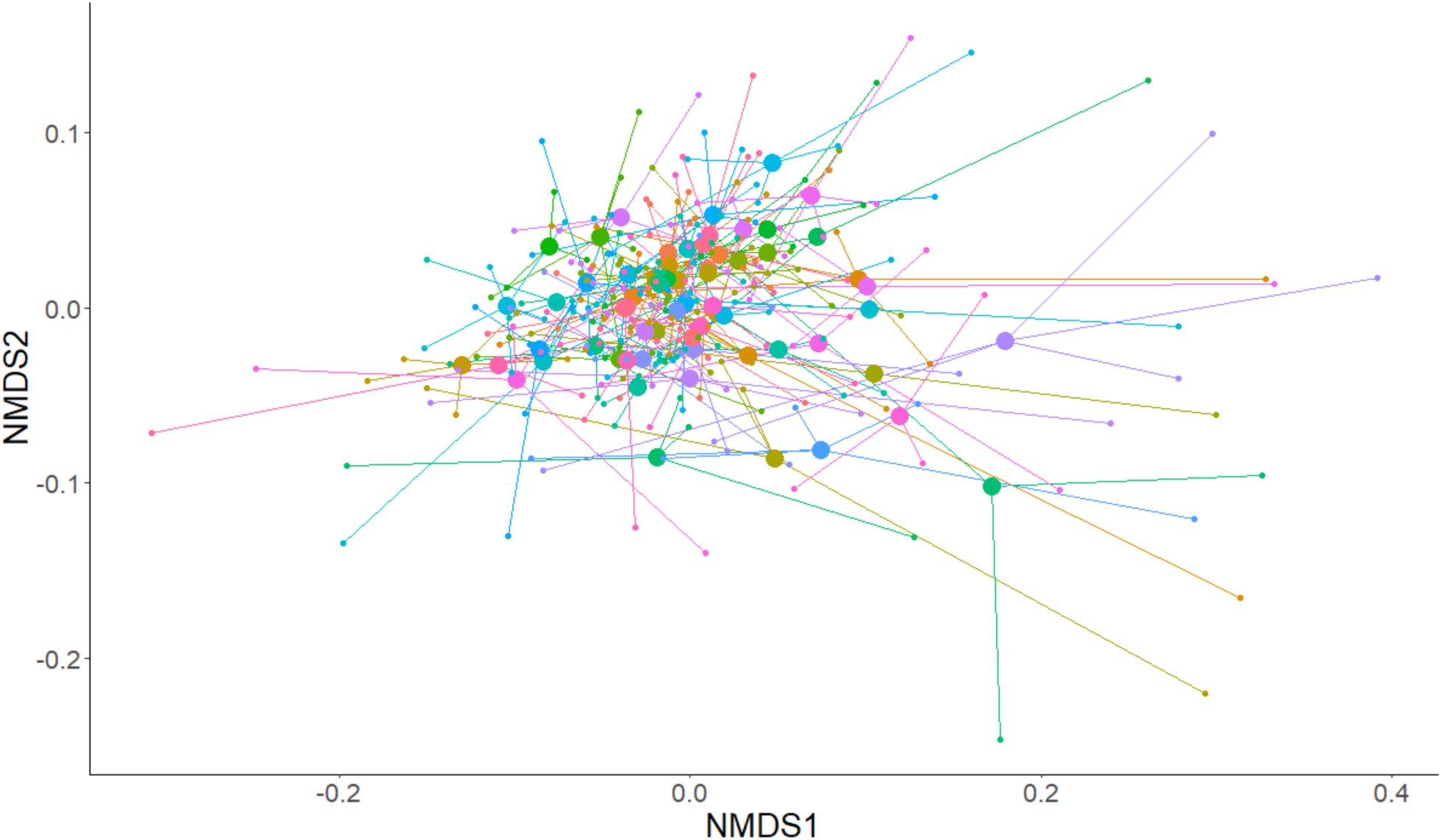
Non-metric multidimensional scaling (nMDS) ordination plot showing the variation among individual honeybee gut bacterial communities, and among the mean colony bacterial communities, across the North West of England. The large circles are the centroid bacterial community for each colony. Each centroid is connected by lines to each of the individuals from that colony, represented by the smaller circles.

### Do colonies that are closer in space have more similar bacterial communities?

We found no evidence that colonies that were closer together geographically possessed more similar gut microbial communities (Mantel test: r=0.020, p=0.336).

### Does land use affect gut bacterial community composition?

There was a marginally significant effect of landscape composition on the pooled gut bacterial community composition of the colonies at the 5km scale (Partial Mantel test: r=0.095, p=0.057), and a trend toward an effect of landscape composition at the 10-km scale (Partial Mantel test: r=0.073, p=0.087). There was no significant effect of landscape composition on pooled colony gut microbial composition at the 500-m (Partial Mantel test: r=0.041, p=0.234) or the 1.5-km scales (Partial Mantel test: r=0.072, p=0.109). The proportion of urban land surrounding a colony did not predict overall composition of the gut microbiota at the colony level (PERMANOVA: R^2^=0.018; df=1, 62; p=0.325).

### Do genetic diversity and host genotype affect gut bacterial community composition?

There were 81 alleles across the five microsatellite loci we studied, with the number of alleles per locus ranging from nine to 22 (Supplementary Table S4). Only 0.14% of the data were null alleles. Of 293 individual bees, only 14 did not have unique genotypes. Thus, the five loci were enough to discriminate among most individuals. None of the loci were in Hardy-Weinberg equilibrium (p<0.001), which is not surprising given that honeybees are a managed species, and the samples include a large number of related individuals. The genetic distances among individuals are shown in Supplementary Figure S2, and the allele richness and heterozygosity data for each colony is reported in Supplementary Table S5. Individual genotype did not influence the gut bacterial community composition of individual bees (Mantel test: r=-0.042, p=0.928), and colony genotype did not influence colony-level gut bacterial community composition (Mantel test: r=-0.128, p=0.967).

### What factors predict alpha diversity in the gut microbiota at the colony and individual level?

At the colony level, we found a marginally significant relationship between the species richness of the gut bacterial community and the proportion of urban land surrounding the colony (p = 0.062; Table 1). The least urban colonies had approximately 3.7 more species in their pooled gut microbiota than the most urban colonies. We found a marginally significant relationship (p = 0.095) between the proportion of alleles shared between pairs of colony-mates and the biodiversity of the gut microbiome at the colony level for q = 1, and a non-significant trend in the same direction (p = 0.101) for q = 2. In both cases, colonies with lower proportions of shared alleles (*i.e*., higher genetic diversity) had more diverse gut bacterial communities. At the individual level, bees from colonies with fewer alleles shared between pairs of workers (i.e. higher genetic diversity) had greater gut bacterial biodiversity when q = 1 (p = 0.040) and when q = 2 (p = 0.040) (Fig 3, Table 2).

**Figure 3.**
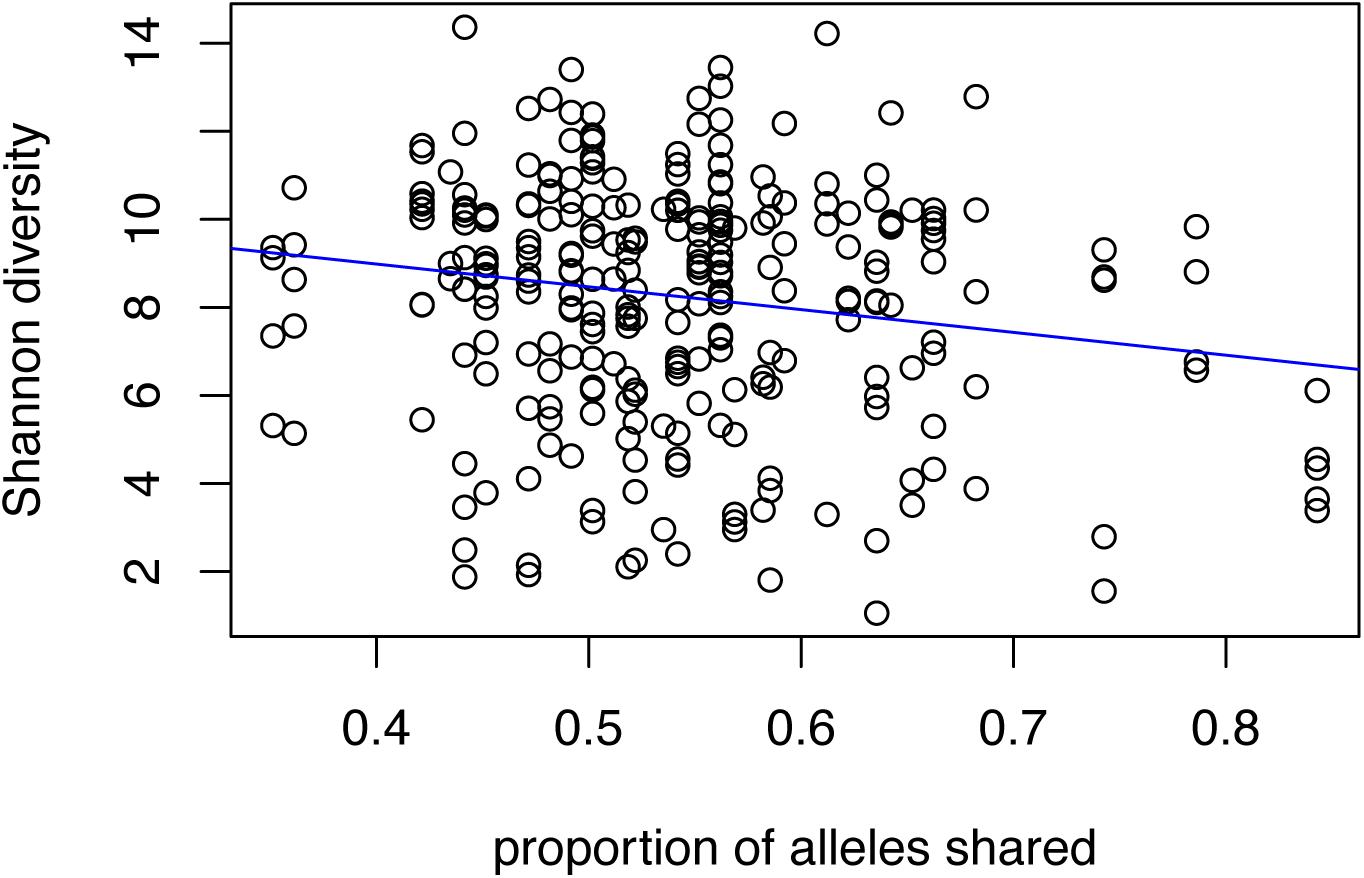
Relationship between the proportion of alleles shared among individuals in a colony and the Shannon diversity (Hill number q=1) of those individuals’ gut bacterial communities. Lower shared alleles indicates higher genetic diversity at the colony level. Therefore, in colonies that are more genetically diverse, individuals had more diverse gut bacteria. The regression line (slope = -0.47, p =. 0.040) is the line of best fit to the data, weighted across all fitted models.

**Table 1.** Factors predicting alpha diversity at the colony level. For each Hill number (q), the first number is the effect size scaled to the standard deviation of the predictor in the population, and the second number is the p-value associated with the predicter. • indicates a marginally significant effect. The effect size is the change in Hill number that we would expect for a change of one standard deviation in the predictor. Positive (negative) effects indicate that alpha diversity increases (decreases) as the predictor increases.

|  | q = 0 |  | q = 1 |  | q = 2 |  |
| --- | --- | --- | --- | --- | --- | --- |
|  | effect | p-value | effect | p-value | effect | p-value |
| proportion of alleles shared between pairs of individuals | -0.33 | 0.486 | -0.51 | <u>0.095</u>• | -0.436 | 0.102 |
| mean heterozygosity | 0.53 | 0.266 | -0.06 | 0.852 | -0.10 | 0.686 |
| landscape diversity | -0.13 | 0.792 | 0.09 | 0.867 | 0.12 | 0.741 |
| proportion of urban land | -0.92 | <u>0.062</u>• | -0.37 | 0.274 | -0.20 | 0.477 |
| northing | -0.22 | 0.667 | -0.23 | 0.496 | -0.13 | 0.642 |
| easting | 0.49 | 0.364 | 0.47 | 0.170 | 0.30 | 0.322 |

**Table 2.**
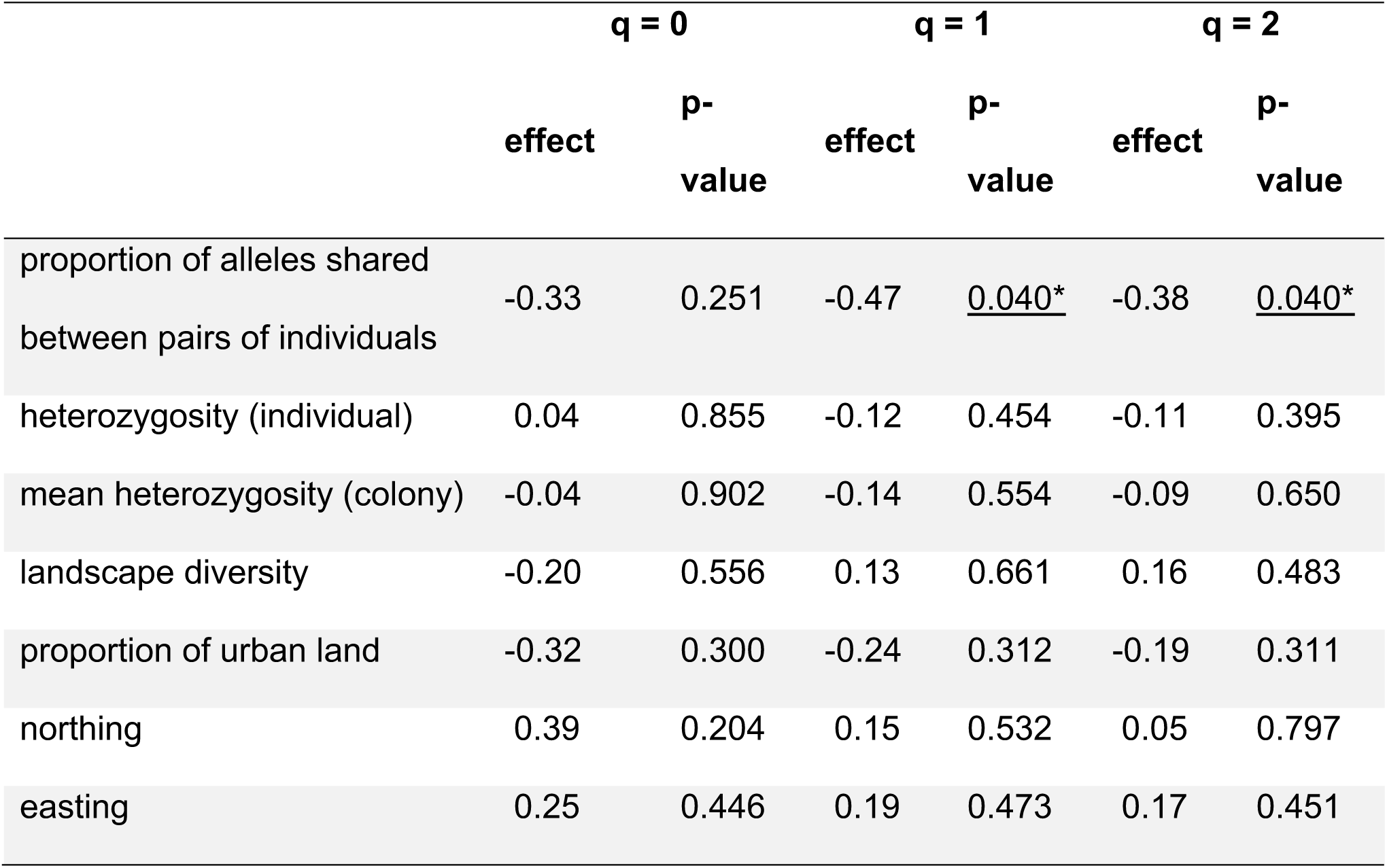
Factors predicting alpha diversity at the individual level. For each Hill number (q), the first number is the effect size scaled to the standard deviation of the predictor in the population, and the second number is the p-value associated with the predictor. * indicates a significant effect. The effect size is the change in the Hill number that we would expect for a change of one standard deviation in the predictor. Positive (negative) effects indicate that alpha diversity increases (decreases) as the predictor increases.

|  | q = 0 |  | q = 1 |  | q = 2 |  |
| --- | --- | --- | --- | --- | --- | --- |
|  | effect | p-value | effect | p-value | effect | p-value |
| proportion of alleles shared between pairs of individuals | -0.33 | 0.251 | -0.47 | <u>0.040*</u> | -0.38 | <u>0.040*</u> |
| heterozygosity (individual) | 0.04 | 0.855 | -0.12 | 0.454 | -0.11 | 0.395 |
| mean heterozygosity (colony) | -0.04 | 0.902 | -0.14 | 0.554 | -0.09 | 0.650 |
| landscape diversity | -0.20 | 0.556 | 0.13 | 0.661 | 0.16 | 0.483 |
| proportion of urban land | -0.32 | 0.300 | -0.24 | 0.312 | -0.19 | 0.311 |
| northing | 0.39 | 0.204 | 0.15 | 0.532 | 0.05 | 0.797 |
| easting | 0.25 | 0.446 | 0.19 | 0.473 | 0.17 | 0.451 |

For an overview of all of the tests and results undertaken see Table 3.

**Table 3.**
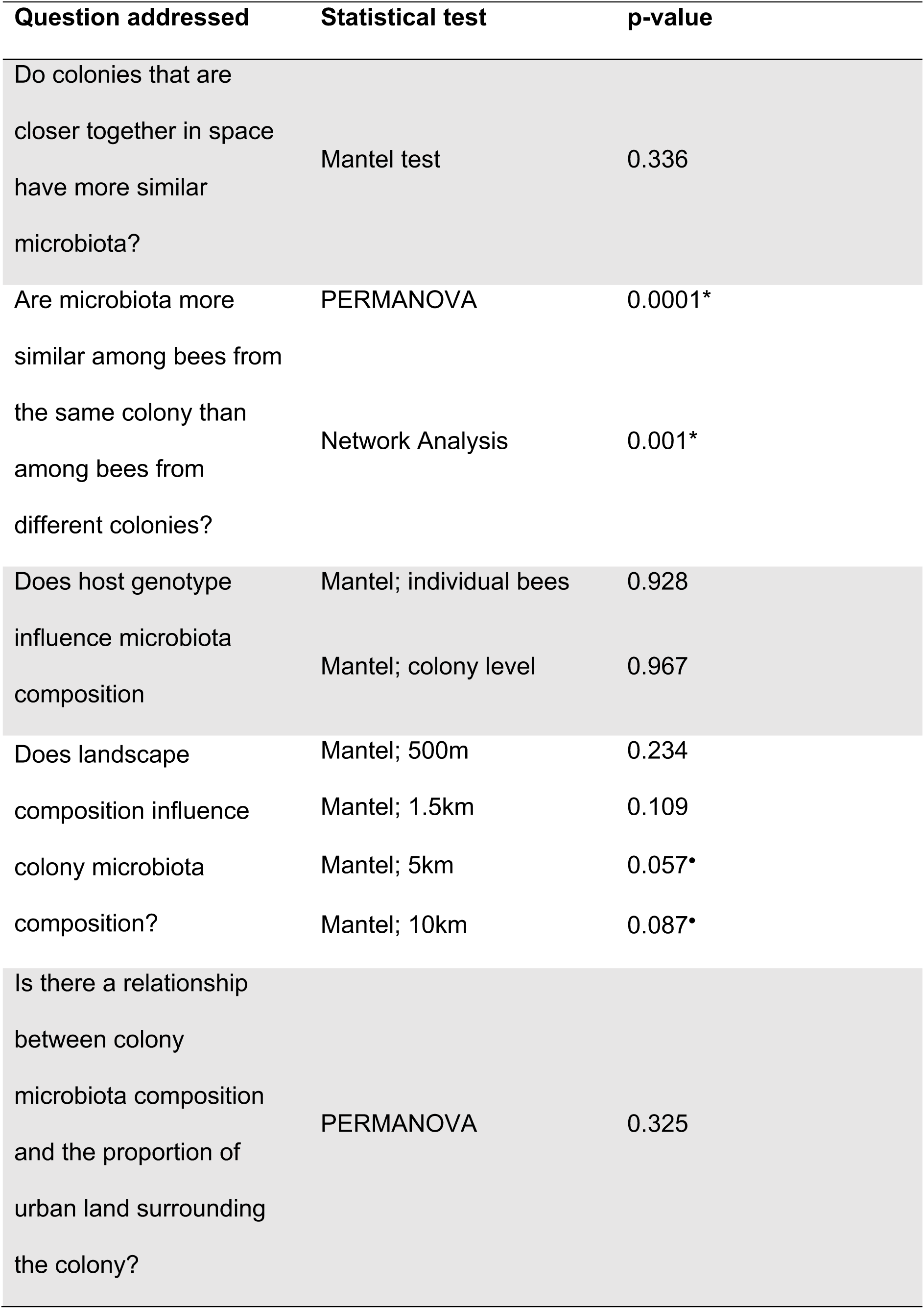
The question addressed, the statistical test used and resulting p-value for the statistical tests performed in this study.

## Discussion

This study reports a detailed investigation of honeybee gut bacterial communities across a geographic region. A few core bacterial phylotypes dominated the communities in our study, and each phylotype was in turn dominated by just one or two ASVs. There was structuring of the microbiota by colony, but an individual’s colony membership could not be predicted from its microbiota composition. We found weak evidence that more urban colonies had less diverse gut bacterial communities, and that colonies from more similar environments had more similar gut bacterial communities. We found stronger evidence that individual bees had greater gut bacterial diversity when their colonies had greater genetic diversity. This last result is particularly interesting, because it means that, in honeybees, the genotypes of individuals predict the bacterial communities of other individuals in the population. To our knowledge, this is the first time this has been shown for any species.

### Honeybee gut bacterial community composition in the North West of England

Our study provides further evidence that the honeybee gut bacterial community is dominated by a core set of phylotypes that have been found consistently across landscapes and continents (Jeyaprakash *et al*., 2003; Mohr and Tebbe, 2006; Babendreier *et al*., 2007; Cox-Foster *et al*., 2007; Ahn *et al*., 2012; Sabree *et al*., 2012; Disayathanoowat *et al*., 2012; Martinson *et al*., 2012; Moran *et al*., 2012; Corby-Harris *et al.,* 2014a; Kwong *et al*., 2017). The only absentee from the literature-defined core in our study was *Parasaccharibacter apium*. The absence of *P. apium* is not surprising as it is generally not found in worker guts, and instead can be found in worker hypopharyngeal glands, the guts of mature queens, larvae and the colony environment (Corby-Harris *et al*., 2016). A lack of *P. apium* in our data suggests our methods successfully limited contamination from bacteria in the colony environment and the worker crop.

The literature-defined core phylotypes comprised a similar proportion of the honeybee gut bacterial community in this study and in previous studies (Moran, 2015). Furthermore, *Lactobacillus* Firm 5 and *Gilliamella apicola* were the most abundant phylotypes in the honeybee gut here, and both have been found to dominate in other studies (Cox-Foster *et al*., 2007; Moran *et al*., 2012). Therefore, our results suggest that the bacterial communities of honeybee guts in North West England are similar to those seen in other places.

Within each core phylotype there were multiple ASVs, but most of the reads within each belonged to just one or two ASVs. Previous studies have found multiple strains of a phylotype within a single individual, with one strain often dominating (Moran *et al*., 2012). The amplicon length used in this study precludes the discrimination of different strains, but the study extends the conclusion of previous studies to show that the same ASVs dominate not just within individuals, but across the region, among different host genotypes, and in bees exposed to different land use types.

### Does land use affect gut bacterial community composition?

In the 5 km and 10 km buffer zones, there was a trend for landscapes with more similar land use composition to harbour colonies with more similar gut bacterial communities. These results are consistent with a small body of work that links honeybee microbiota to land use. Jones *et al*. (2018) found small differences in the gut microbiota of colonies placed near oil seed rape fields and those placed further away. Donkersley *et al*. (2018) found that land use surrounding colonies predicts the species richness of the bee bread microbiota. In contrast, in three bumblebee species, Cariveau *et al*. (2014) found no difference in the microbiota of individuals collected at an active or at an abandoned cranberry farm.

We found a weak trend linking the diversity of the honeybee gut bacterial community to urbanisation. In more urban environments, gut bacterial communities had lower species richness. Among synanthropic bird species, similar patterns have been reported in house sparrows (*Passer domesticus*; Teyssier *et al*., 2018) in Belgium and herring gulls (*Larus argentatus*; Fuirst *et al*., 2018) in New York, but the opposite pattern was reported in white-crowned sparrows (*Zonotrichia leucophrys*; Phillips *et al*., 2018) in California. Bosmans *et al*. (2018) reported higher gut bacterial diversity in bumblebee (*Bombus terrestris*) queens from two forested sites than from three urban sites in Belgium, but with only five sites sampled it was impossible to attribute differences with confidence to urbanisation.

### Do genetic diversity and host genotype affect gut bacterial community composition?

We found that individual bees from more genetically diverse colonies had more diverse gut bacterial communities. In a study in which colony genetic diversity was experimentally manipulated, Mattila *et al*. (2012) found that colonies with more genetic diversity had more diverse bacterial communities. Our results advance those of Mattila and colleagues in two important ways. First, the genetic diversity in our study was not manipulated, so our results show that naturally occurring differences in genetic diversity are sufficient to predict the diversity of gut bacterial communities. Second, the differences in bacterial diversity uncovered by Matilla and colleagues appeared at the level of the colony. Such differences could occur in two non-exclusive ways. One possibility is that bees with different genotypes could host different bacterial communities. Individual bees in genetically diverse colonies might not host more diverse communities themselves, but if each host a different set of bacteria, then the pooled community at the colony level would be more diverse. The other possibility is that, in more genetically diverse colonies, individual bees host more diverse bacterial communities. In our study, the latter explanation was true. This shows that the genotype of each individual in the colony predicts the microbiome of other individuals.

Our results do not show that the genetic diversity at the colony level *causes* diversity in the gut bacterial community. For example, it might be true that colonies in areas with a higher density of colonies have more potential for outbreeding, and also have more potential to exchange microbiota. If this is true, then the relationship between genetic diversity and gut bacterial community diversity might be driven entirely by the environment. We believe the results of Matilla *et al*. (2018) make this explanation unlikely, but additional work will be needed to demonstrate this conclusively.

## Conclusion

This study provides a comprehensive analysis of spatial variation in honeybee gut bacterial community at the regional scale, and provides new evidence for effects of land use and host genetic diversity on the gut microbiota. Given the large number of hypotheses we studied (Supplementary Tables S6-S8), we should expect at least some false positive results, and future work should seek to replicate the patterns we report. Our results provide the foundation for this and other work, and advance our general understanding of the honeybee microbiota.

## Acknowledgments

CB was funded by a UKRI Biotechnology and Biological Sciences Research Council (BBSRC) DTP scholarship (BB/M011208/1). LRV was funded by a Daphne Jackson Trust Fellowship through BBSRC and NERC. Many thanks to all the beekeepers for volunteering to be part of the study and to the local BBKA groups for facilitating connections.

## Author contributions

JKR, WM, LRV and CB conceived and designed the study; LV recruited beekeepers and collected samples; CB and REA performed the molecular laboratory work; CB conducted bioinformatics; CB, TRG and JKR performed data analysis; CB, TRG and JKR wrote the manuscript. All authors gave final approval for publication.

