## Supplementary data files for "The genetic diversity of honeybee colonies predicts the gut bacterial diversity of individual colony members"

---

[1]Faculty of Science and Engineering, University of Manchester, Manchester, UK, M13 9PT

[2]Ecology and Environment Research Centre, Department of Natural Sciences, Manchester Metropolitan University, Manchester, UK, M1 5GD

[3]School of Science, Engineering and Environment, University of Salford, Salford, UK, M5 4WT

[4]Faculty of Biology Medicine and Health, Lydia Becker Institute of Immunology and Inflammation, University of Manchester, Manchester, UK, M13 9PT

[5]Miltenyi Biotec, Bergisch Gladbach, Germany

[6]Department of Infectious Diseases, Medical Microbiology and Hygiene, University of Heidelberg, Heidelberg, Germany

[7]Translational Lung Research Centre (TLRC), Heidelberg, Germany

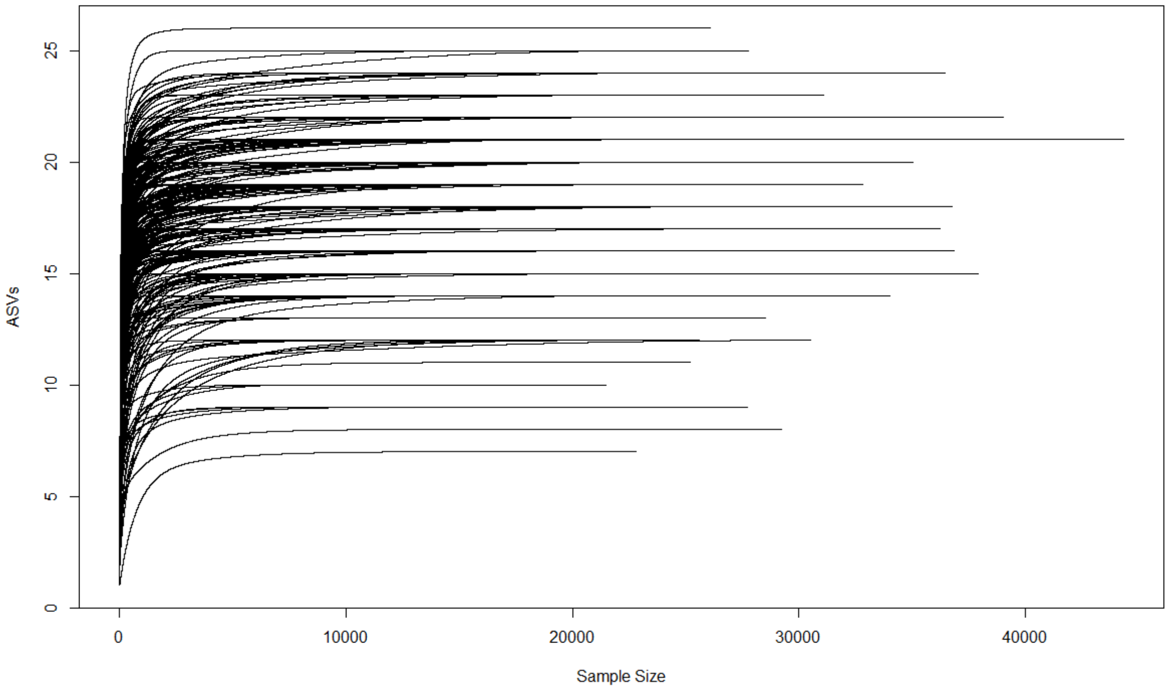


**Figure S1.** Rarefaction curves for each of the 293 honeybee gut samples sequenced using the 515F and 806R universal primers to amplify the V4 region of the 16S rRNA gene on the MiSeq platform (Illumina v2 chemistry).

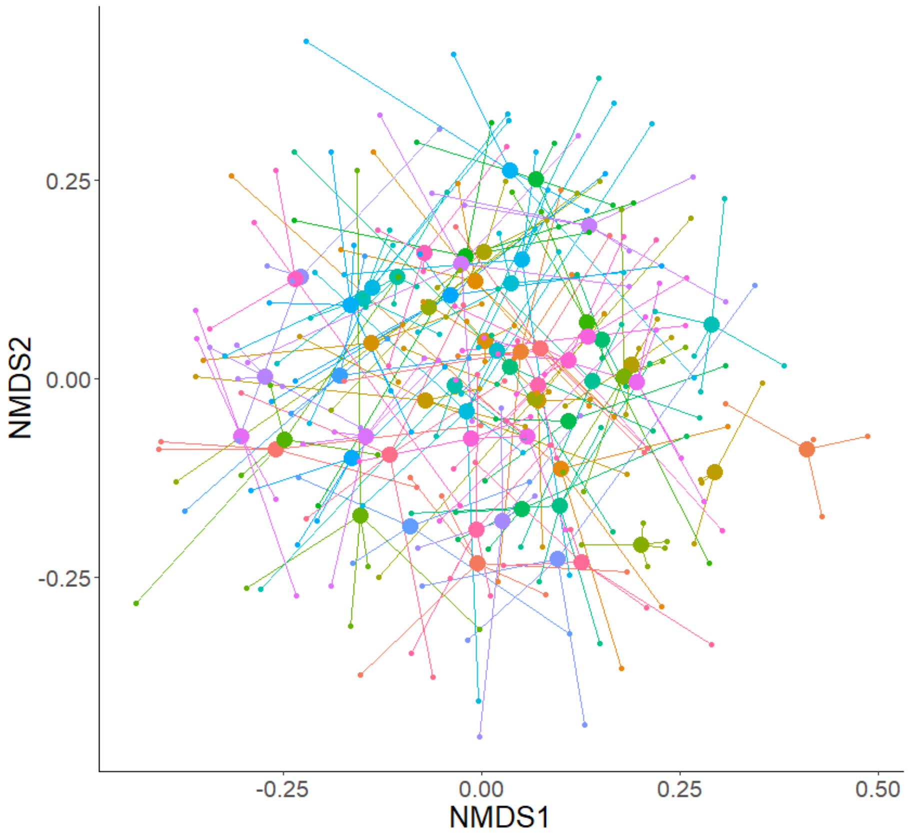


**Figure S2.** Non-metric Multidimensional Scaling (nMDS) ordination of Roger's genetic distances among individual honeybees. Each small data point represents an individual bee, with lines connecting individuals to their colony centroid (larger circles).

**Table S1.** Primer pairs used to amplify the five microsatellite loci (from Evans *et al.* 2013).

| Primer Name | Sequence |
|---|---|
| A113-F-(FAM) | CTC GAA TCG TGG CGT CC |
| A113-R | CCT GTA TTT TGC AAC CTC GC |
| A007-F-(TAMRA) | GTT AGT GCC CTC CTC TTG C |
| A007-R | CCC TTC CTC TTT CAT CTT CC |
| AP043-F-(JOE) | GGC GTG CAC AGC TTA TTC C |
| AP043-R | CGA AGG TGG TTT CAG GCC |
| AP055-F-(ROX) | GAT CAC TTC GTT TCA ACC GT |
| AP055-R | CAT TCG GTA TGG TAC GAC CT |
| B124-F-(FAM) | GCA ACA GGT CGG GTT AGA G |
| B124-R | CAG GAT AGG GTA GGT AAG CAG |

**Table S2.** Summary of the abundances of the core gut bacterial taxa found from 63 hives in NW England including: the total number of reads of all ASVs associated with a species cluster; the total number of ASVs belonging to each core species cluster (Number of Variants); the percentage of reads of a species cluster that belong to the most abundant ASV of that cluster (% Most Abundant Variant); the percentage abundance of that species cluster in the region as a whole; and the percentage abundance of the species cluster within only the top 20 most abundant ASVs.

| Taxa | Number of Reads | Number of Variants | % Most Abundant Variant | Overall Abundance (%) | Abundance in Top 20 ASVs (%) |
|---|---|---|---|---|---|
| *Gilliamella* | 1383612 | 13 | 97.6 | 20.1 | 20.7 |
| *Frischella* | 479766 | 8 | 75.5 | 7.0 | 7.0 |
| *Snodgrassella* | 750352 | 5 | 77.4 | 10.9 | 11.5 |
| *Lactobacillus* Firm 4 | 511096 | 14 | 37.5 | 7.4 | 6.9 |
| *Lactobacillus* Firm 5 | 1965934 | 25 | 28.3 | 28.5 | 28.6 |
| *Bifidobacteria* | 832217 | 5 | 94.4 | 12.1 | 12.7 |
| *Bartonella* | 568797 | 3 | 93.9 | 8.3 | 8.2 |
| *Commensalibacter* | 266042 | 8 | 66.4 | 3.9 | 3.5 |

**Table S3.** The prevalence of each of the core species clusters across individual samples and for samples pooled at the colony level. Prevalence is reported as the percentage of individuals or colonies that possess at least one ASV from the species cluster.

| Species Cluster | Presence in Individuals/ % | Presence in Colonies/ % |
|---|---|---|
| *Gilliamella* | 99.7 | 100 |
| *Frischella* | 92.5 | 100 |
| *Snodgrassella* | 98.3 | 100 |
| *Lactobacillus* Frim 4 | 93.9 | 100 |
| *Lactobacillus* Firm 5 | 100 | 100 |
| *Bifidobacteria* | 98.3 | 100 |
| *Bartonella* | 90.8 | 100 |
| *Commensalibacter* | 83.6 | 100 |

**Table S4.** Number of alleles, expected ($H_e$) and observed ($H_o$) Heterozygosity for each of the five microsatellite loci used for genotyping in this study.

| **Locus** | **Number of alleles** | **$H_e$** | **$H_o$** |
|---|---|---|---|
| A113 | 16 | 0.69 | 0.66 |
| AP043 | 9 | 0.76 | 0.70 |
| AP055 | 16 | 0.80 | 0.81 |
| A007 | 22 | 0.78 | 0.77 |
| B124 | 18 | 0.87 | 0.85 |

**Table S5.** The total number of alleles, individual Heterozygosity and mean Heterozygosity ($H_s$) across all loci for each of the 63 honeybee colonies sampled in NW England.

| Colony ID | Number of alleles | Individual heterozygosity | | | | | |
|---|---|---|---|---|---|---|---|
| | | **A113** | **AP043** | **AP055** | **A007** | **B124** | **$H_s$** |
| ADW | 17 | 0.37 | 0.35 | 0.63 | 0.8 | 0.75 | 0.58 |
| AGE | 14 | 0.37 | 0.45 | 0.68 | 0.72 | 0.55 | 0.554 |
| AND | 14 | 0.71 | 0.25 | 0.58 | 0.25 | 0.67 | 0.492 |
| ANDZ | 15 | 0.6 | 0.5 | 0.6 | 0.68 | 0.75 | 0.626 |
| AR | 22 | 0.88 | 0.5 | 0.83 | 0.83 | 0.67 | 0.742 |
| ARZ | 22 | 0.68 | 0.8 | 0.72 | 0.75 | 0.82 | 0.754 |
| AT | 17 | 0.5 | 0.68 | 0.7 | 0.63 | 0.68 | 0.638 |
| BIC | 17 | 0.58 | 0.58 | 0.46 | 0.83 | 0.83 | 0.656 |
| BPO | 17 | 0.65 | 0.7 | 0.6 | 0.73 | 0.68 | 0.672 |
| CAT | 16 | 0.68 | 0.7 | 0.75 | 0.8 | 0.45 | 0.676 |
| CGH | 12 | 0.5 | 0.42 | 0.5 | 0.62 | 0.62 | 0.532 |
| CGR | 19 | 0.53 | 0.6 | 0.75 | 0.53 | 0.78 | 0.638 |
| CHA | 15 | 0.42 | 0.42 | 0.62 | 0.71 | 0.71 | 0.576 |
| CK | 14 | 0.75 | 0.75 | 0.75 | 0.75 | 0.5 | 0.7 |
| CWI | 15 | 0.6 | 0.78 | 0.75 | 0.35 | 0.68 | 0.632 |
| DEB | 14 | 0.8 | 0.35 | 0.65 | 0.5 | 0.6 | 0.58 |
| DEL | 15 | 0.35 | 0.4 | 0.75 | 0.5 | 0.68 | 0.536 |
| DJ | 17 | 0.2 | 0.75 | 0.6 | 0.78 | 0.7 | 0.606 |
| DLY | 15 | 0.33 | 0.67 | 0.75 | 0.92 | 0.67 | 0.668 |

| | | Individual heterozygosity | | | | | |
|---|---|---|---|---|---|---|---|
| **Colony** | **Number of alleles** | **A113** | **AP043** | **AP055** | **A007** | **B124** | **$H_s$** |
| EAD | 18 | 0.5 | 0.6 | 0.78 | 0.72 | 0.75 | 0.67 |
| EFL | 16 | 0.46 | 0.46 | 0.75 | 0.71 | 0.62 | 0.6 |
| GB | 14 | 0.5 | 0.58 | 0.5 | 0.5 | 0.92 | 0.6 |
| HAR | 16 | 0.75 | 0.67 | 0.75 | 0.42 | 0.79 | 0.676 |
| HMO | 12 | 0.67 | 0.75 | 0.5 | 0 | 0.67 | 0.518 |
| HSS | 18 | 0.7 | 0.55 | 0.65 | 0.7 | 0.79 | 0.678 |
| IL | 16 | 0.8 | 0.65 | 0.5 | 0.7 | 0.68 | 0.666 |
| IMO | 17 | 0.65 | 0.68 | 0.68 | 0.53 | 0.78 | 0.664 |
| IRLO | 15 | 0.5 | 0.5 | 0.62 | 0.7 | 0.75 | 0.614 |
| JA | 14 | 0.67 | 0.71 | 0.25 | 0.67 | 0.67 | 0.594 |
| JCH | 15 | 0.2 | 0.65 | 0.62 | 0.37 | 0.75 | 0.518 |
| JENE | 18 | 0.58 | 0.53 | 0.7 | 0.72 | 0.65 | 0.636 |
| JGPH | 15 | 0.5 | 0.75 | 0.58 | 0.67 | 0.75 | 0.65 |
| JHW | 17 | 0.45 | 0.65 | 0.65 | 0.75 | 0.73 | 0.646 |
| JL | 15 | 0.7 | 0.55 | 0.35 | 0.85 | 0.65 | 0.62 |
| JN | 16 | 0.6 | 0.5 | 0.35 | 0.7 | 0.8 | 0.59 |
| KIN | 20 | 0.53 | 0.78 | 0.75 | 0.72 | 0.72 | 0.7 |
| KK | 16 | 0.42 | 0.62 | 0.46 | 0.71 | 0.62 | 0.566 |
| KTH | 20 | 0.82 | 0.7 | 0.72 | 0.75 | 0.63 | 0.724 |
| LIS | 17 | 0.65 | 0.6 | 0.6 | 0.75 | 0.7 | 0.66 |
| LWA | 17 | 0.6 | 0.75 | 0.45 | 0.63 | 0.8 | 0.646 |
| MAD | 17 | 0.45 | 0.8 | 0.68 | 0.62 | 0.55 | 0.62 |
| MAN | 19 | 0.55 | 0.6 | 0.8 | 0.68 | 0.82 | 0.69 |

| | | Individual heterozygosity | | | | | |
|---|---|---|---|---|---|---|---|
| **Colony** | **Number of alleles** | **A113** | **AP043** | **AP055** | **A007** | **B124** | **$H_s$** |
| MB | 16 | 0.79 | 0.67 | 0.42 | 0.79 | 0.62 | 0.658 |
| MPO | 13 | 0.55 | 0.6 | 0.45 | 0.5 | 0.58 | 0.536 |
| MST | 20 | 0.82 | 0.7 | 0.75 | 0.62 | 0.75 | 0.728 |
| ND | 13 | 0 | 0.35 | 0.6 | 0.6 | 0.8 | 0.47 |
| NW | 18 | 0.37 | 0.68 | 0.78 | 0.6 | 0.78 | 0.642 |
| PBO | 18 | 0.62 | 0.7 | 0.4 | 0.75 | 0.85 | 0.664 |
| PGS | 10 | 0.35 | 0.5 | 0.5 | 0.5 | 0.35 | 0.44 |
| PL | 15 | 0 | 0.71 | 0.67 | 0.58 | 0.75 | 0.542 |
| PMA | 19 | 0.5 | 0.63 | 0.72 | 0.68 | 0.85 | 0.676 |
| REN | 15 | 0.71 | 0.5 | 0.67 | 0.42 | 0.88 | 0.636 |
| RW | 17 | 0.58 | 0.62 | 0.71 | 0.71 | 0.83 | 0.69 |
| SH | 15 | 0.67 | 0.58 | 0.67 | 0.67 | 0.67 | 0.652 |
| SIW | 18 | 0.58 | 0.68 | 0.7 | 0.78 | 0.75 | 0.698 |
| SSP | 19 | 0.78 | 0.65 | 0.8 | 0.6 | 0.68 | 0.702 |
| TAY | 15 | 0.67 | 0.67 | 0.75 | 0.58 | 0.58 | 0.65 |
| TEM | 17 | 0.5 | 0.78 | 0.7 | 0.6 | 0.72 | 0.66 |
| TSE | 23 | 0.8 | 0.85 | 0.78 | 0.72 | 0.88 | 0.806 |
| VCU | 19 | 0.7 | 0.45 | 0.58 | 0.75 | 0.72 | 0.64 |
| VHA | 22 | 0.5 | 0.78 | 0.63 | 0.82 | 0.82 | 0.71 |
| WIL | 21 | 0.8 | 0.68 | 0.8 | 0.53 | 0.88 | 0.738 |
